# From reporters to endogenous genes: the impact of the first five codons on translation efficiency in *Escherichia coli*

**DOI:** 10.1101/699900

**Authors:** Mariana H. Moreira, Géssica C. Barros, Rodrigo D. Requião, Silvana Rossetto, Tatiana Domitrovic, Fernando L. Palhano

**Author notes:** These authors contributed equally (Mariana H. Moreira, Géssica C. Barros, Rodrigo D. Requião).

## Abstract

Translation initiation is a critical step in the regulation of protein synthesis, and it is subjected to different control mechanisms, such as 5’ UTR secondary structure and initiation codon context, that can influence the rates at which initiation and consequentially translation occur. For some genes, translation elongation also affects the rate of protein synthesis. With a GFP library containing nearly all possible combinations of nucleotides from the 3^rd^ to the 5^th^ codon positions in the protein coding region of the mRNA, it was previously demonstrated that some nucleotide combinations increased GFP expression up to four orders of magnitude. While it is clear that the codon region from positions 3 to 5 can influence protein expression levels of artificial constructs, its impact on endogenous proteins is still unknown. Through bioinformatics analysis, we identified the nucleotide combinations of the GFP library in *Escherichia coli* genes and examined the correlation between the expected levels of translation according to the GFP data with the experimental measures of protein expression. We observed that *E. coli* genes were enriched with the nucleotide compositions that enhanced protein expression in the GFP library, but surprisingly, it seemed to affect the translation efficiency only marginally. Nevertheless, our data indicate that different enterobacteria present similar nucleotide composition enrichment as *E. coli*, suggesting an evolutionary pressure towards the conservation of short translational enhancer sequences.

## INTRODUCTION

Protein synthesis is the most energetically costly process for a bacteria cell^1^. Therefore, to ensure that the benefit due to the expression of a given gene exceeds or equals the costs of its production, several regulatory mechanisms operate not only at the level of transcription but also at the translation step to optimize the final protein output. It has long been known that the yield of proteins produced per mRNA molecule, or translation efficiency (TE), can be greatly affected by structural elements and sequence motifs at the mRNA 5’UTR^2^. For example, both the nucleotide composition and structural context of the Shine-Dalgarno sequence determine protein translation initiation and translation efficiency in bacteria^3^. More recently, it has been demonstrated that the expression levels of some proteins are also regulated by translation elongation rates^4,5^. Different studies agree that in addition to initiation, early elongation events (first 10-30 amino acid-coding nucleotides) are relevant for TE^6^. The recognized regulatory factors are tRNA abundance for a given codon, nucleotide composition, mRNA structure, and the nascent polypeptide sequence. However, because these variants are not completely independent, it is difficult to pinpoint the major determinant for the observed effect, generating conflicting models for translational efficiency regulation^7–11^.

Heterologous reporter libraries designed to test the influence of a specific region of a transcript on TE became a powerful approach to identify regulatory elements of translation. The constructs have invariable promoters and 5’ UTRs, which homogenize transcriptional levels and initiation rates. Moreover, by fixing first two amino acids of a protein, posttranslational protein degradation by the N-end rule pathway is also controlled. Hence, TE can be estimated directly from protein expression levels. Very recently, Djuranovic’s group reported the effect of the five first codons on translation efficiency in *E. coli* ^12^. The authors generated the most comprehensive GFP library available so far, with virtually all-possible nucleotide compositions for codons 3 to 5 (262,144). Recombinant cells were FACS-sorted into 5 different groups according to their fluorescence levels, and the plasmids were sequenced to quantify the occurrence of specific sequences in each group. In this manner, each GFP construct was scored according to the resulting fluorescence intensity (GFP score). The authors also derived a nucleotide and an amino acid motif that was more common among highly expressed constructs. Interestingly, some nucleotide compositions enhanced the magnitude of protein abundance up to 4 orders in relation to the original construct, without interfering with the mRNA levels. In other words, the nucleotide composition of the beginning of the protein coding sequence drastically affected the final protein output. The authors showed that this phenomenon could be observed with other protein reporters, independent of the expression vector used, or *in vitro* and *in vivo* expression conditions. While this kind of study has an unquestionable value for optimizing the production of heterologous constructs, it is still debated whether this experimental setting is too artificial, so that, in a genomic context, the effect of these putative regulatory motifs would be overruled by stronger determinant factors of protein expression.

In this work, our aim was to verify the biological significance of the GFP score obtained with this GFP library. In other words, we wanted to evaluate whether the elongation regulatory motifs that were identified by high-throughput studies can be used to predict the expression of endogenous *E. coli* genes. Through bioinformatics, we searched the *E. coli* genome for each nucleotide composition of the GFP library, attributing the respective GFP-score to each gene (GFP-predicted score). We observed that the nucleotide compositions that yielded higher GFP-scores were overrepresented in *E. coli* genes when compared to a scrambled version of the genome. This result suggested a preference for sequences that increased translatability at the elongation step. However, the correlation matrix between the GFP-predicted score vs. previously published experimental parameters related to gene expression, such as translation efficiency, protein abundance, mRNA abundance, and mRNA stability, showed no significance (Spearman correlation ranging from −0.08 to 0.13). Nevertheless, when evaluated as a group, we observed that genes with high GFP-predicted scores had higher TE values when compared with the genes with low scores. Moreover, the ribosome occupancy at codon positions 3 to 5 was lower when compared to the average of the genome, suggesting that this short region modulates the speed of translation. Finally, we observed that the bias towards sequences with high GFP-predicted scores was conserved in other enterobacterial species, such as *Klebsiella oxytoca*, and *Enterobacter asburiae*. Additionally, the top scored genes from each of the three enterobacteria had a higher orthology frequency than what would be expected by random chance.

## RESULTS AND DISCUSSION

### Natural occurrence of short elongation-regulatory sequences related to high mRNA translatability identified by high-throughput studies

Previous works indicated that the regulatory effect of early elongation steps on translation efficiency could arise from interactions of the nascent peptide with the ribosomal exit tunnel or from the decoding efficiency of the mRNA nucleotide sequence^13–16^. To further investigate how these factors influence the efficiency of protein synthesis, Verma et al. constructed the most comprehensive eGFP library published so far. They focused on the region surrounding the +10 nucleotide position by introducing 9 random nucleotides after the second codon^12^. The global expression of the eGFP library was FACS-sorted into five bins, normalized by DNA amount and sequenced. A GFP score was calculated to represent the weighted distribution value for each independent sequence over five FACS-sorted bins. By this metric, sequences exclusively associated with bin 1 (lowest expression) would have a score of 1, and sequences exclusively associated with bin 5 (highest expression) would have a score of 5. A GFP score for each of the 259,134 unique sequences was calculated, which covered 98.8% of all possible nucleotide combinations at codon positions 3-5 (4^9^ = 262,144). A cut-off of 10 reads was used, and a Pearson correlation of 0.79 between the replicates was found. For our analyses, the sequences with stop codons were eliminated, and only sequences found in both replicates were used (182,289 sequences, Pearson correlation = 0.70, Figure 1A). A linear fit equation with Q = 1% was used to eliminate the constructs with low reproducibility (29,945 sequences), and the geometrical mean of the replicates was used to score 152,344 unique nucleotide sequences (Figure 1A). The frequency of distribution of these refined GFP-datasets showed a non-Gaussian distribution with a main peak and median score of 3.10 (Figure 1B, green line). We used this refined dataset to screen the sequence variants among the *E. coli* genes and attribute the corresponding GFP score (GFP predicted score) (Figure 1A). First, we analyzed the score at codons 3-5, the same positions used in the GFP library. Using this approach, we could attribute a score for 2,644 genes with 2,500 different sequence combinations, which covered 64% of the *E. coli* genome (Figure 1B, red line). Similar results were obtained by Djuranovic’s group (personal communication). The density histogram obtained with the *E. coli* genome differed from the GFP library, showing a higher median GFP-predicted score of 3.23 and an increased frequency of sequences with scores higher than 4 (Figure 1B). We created a scrambled version of each mRNA sequence maintaining the proportions of all 61 amino acid codons (genomic codon bias) but randomizing the codon choice for each transcript. The distribution obtained with this control data set was similar to the GFP library. (Figure 1B, black line). To investigate whether the genomic distribution was position-specific, we scored the *E. coli* genes using codon positions 9-11. In this case, the distribution was similar to that obtained with the scrambled control and the GFP library (Figure 1B, orange line).

**Figure 1.**
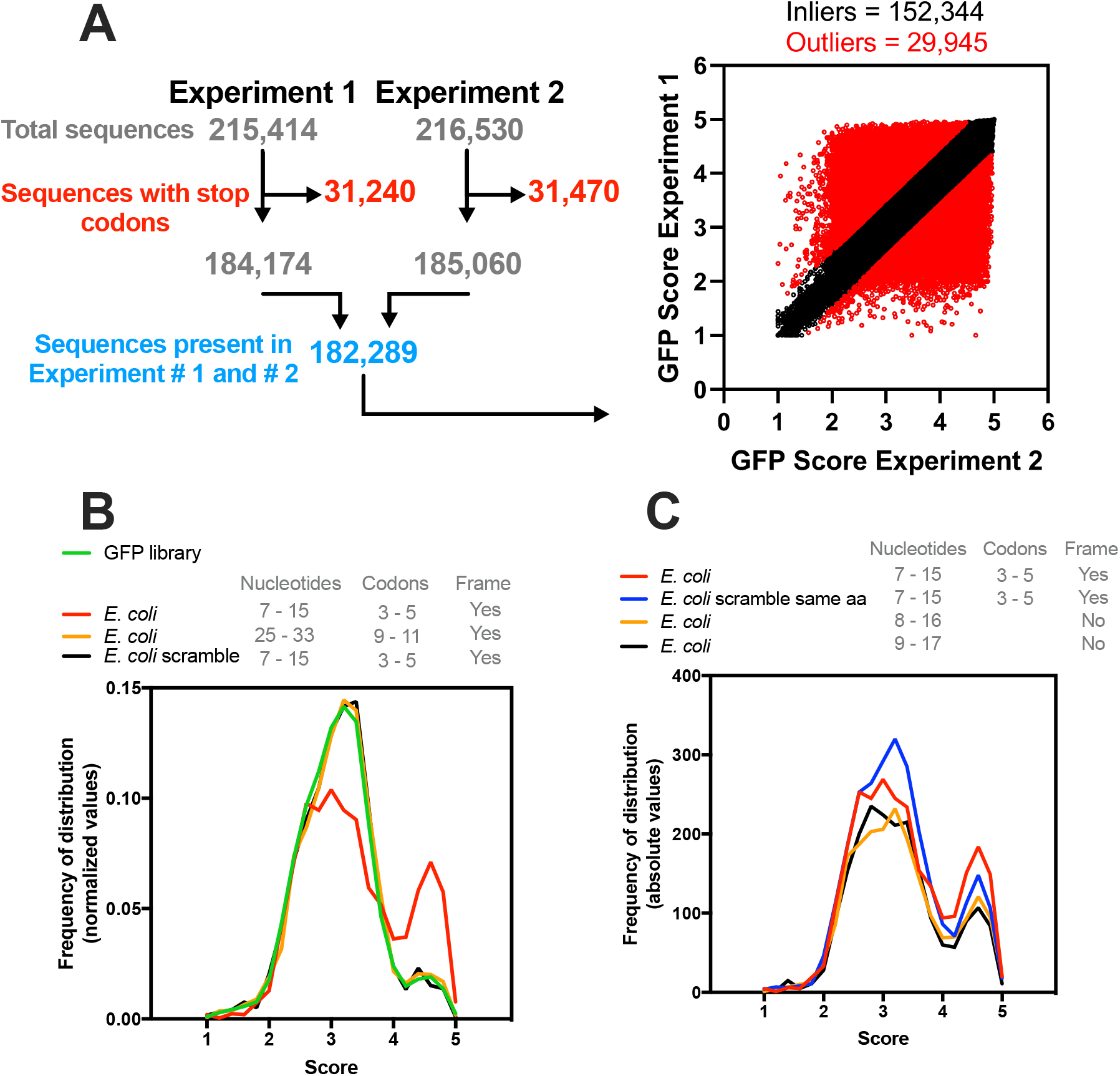
The *E. coli* genome has a bias towards nucleotide composition at codons 3-5. (A) Djuranovic’s group performed a high-throughput screen varying the nucleotide composition at codons 3-5 in GFP. In experiments #1 and #2, 215,414 and 261,530 different compositions were analyzed, respectively, regarding the GFP fluorescence levels. The sequences with a stop codon (31,240 and 31,470 for experiments #1 and #2, respectively) were removed, and only sequences present in both experiments were used (182,289). The outliers (29,945) were defined by setting Q = 1% in the linear regression. We then calculated the average GFP score of the inliers (152,344) from experiments #1 and #2. This list was used in all subsequent bioinformatics experiments. (B) Density histogram of GFP scores for genes identified in *E. coli*. The nucleotide composition at codon positions 3-5 or 9-11 was analyzed. As a control, we used a scrambled genome where the codon proportion was maintained, but their position was randomly changed. Note that only codon positions 3-5 in the real genome possessed a bias towards high GFP scores. (C) The effect of amino acid composition and mRNA sequence on GFP score bias was analyzed. As a control, we used a scrambled genome where the codons were randomly changed, keeping the codon proportion and amino acid sequence of each gene (*E. coli* scramble same aa).

To address the effect of the reading frame on the distribution bias observed for the *E. coli* genome, we shifted the analyzed nucleotide window (+1, 8-16; +2, 9-17). With this modification, fewer genes were ranked with scores > 4, but we still observed a bias towards higher scores, which suggested that both the amino acid and nucleotide sequences were important to the observed distribution. Then, we tested whether the tri-peptide sequence would be enough to generate the same bias observed with the nucleotide sequence. We maintained the proportions of all 61 amino acid codons as well as the final protein sequence but randomized the codon choice for each transcript of *E. coli* (Figure 1C, blue line). We observed only a slight reduction in the bias towards high GFP-predicted scores, confirming that the nascent peptide is important for the translatability of the mRNA. These results were in line with experimental data observed with the GFP constructs and indicated that both the amino acid and nucleotide sequences that confer higher translation efficiency in GFP reporters are more frequent in *E. coli* genes than what would be expected by random chance.

### Influence of short elongation-regulatory sequences identified by high-throughput studies on endogenous gene expression

The GFP score calculated by Verma et al. can be used to estimate the relative expression levels of each eGFP variant in the library. Hence, we asked whether the GFP-predicted scores of endogenous genes (SCORE GENOME) would correlate with the following experimental measurements associated with gene expression levels: TE, determined by ribosome profiling and calculated from the ratio between ribosome-protected footprint read counts and total mRNA read counts for each gene ^17–19^; protein abundance, measured by mass spectrometry ^18,20^; mRNA abundance, determined by RNAseq and microarray ^19,21–23^ and mRNA secondary structure, determined by the extent of chemical accessibility of RNA to dimethyl sulfate and quantified by the Gini index, where the higher the structural content, the higher the Gini value is ^19^. As a control, we used a scrambled *E. coli* genome that maintained the proportions of all 61 amino acid codons but randomized the codon choice for each transcript (SCORE SCRAMBLE).

Three main clusters were found. Cluster 1 was on the isolated branch formed by the negative correlation values obtained with the RNA structure Gini Index and other analyzed parameters (Figure 2A). This was expected, as a high degree of mRNA secondary structure can slowdown translation elongation ^19^. The second cluster was formed by TE, protein abundance, and mRNA levels that had a high positive correlation between them (Figure 2A). SCORE GENOME and SCORE SCRAMBLE formed cluster 3, which had no correlation to each other or with the other parameters analyzed (Spearman correlation ranging from −0.08 to 0.13) (Figure 2A).

**Figure 2.**
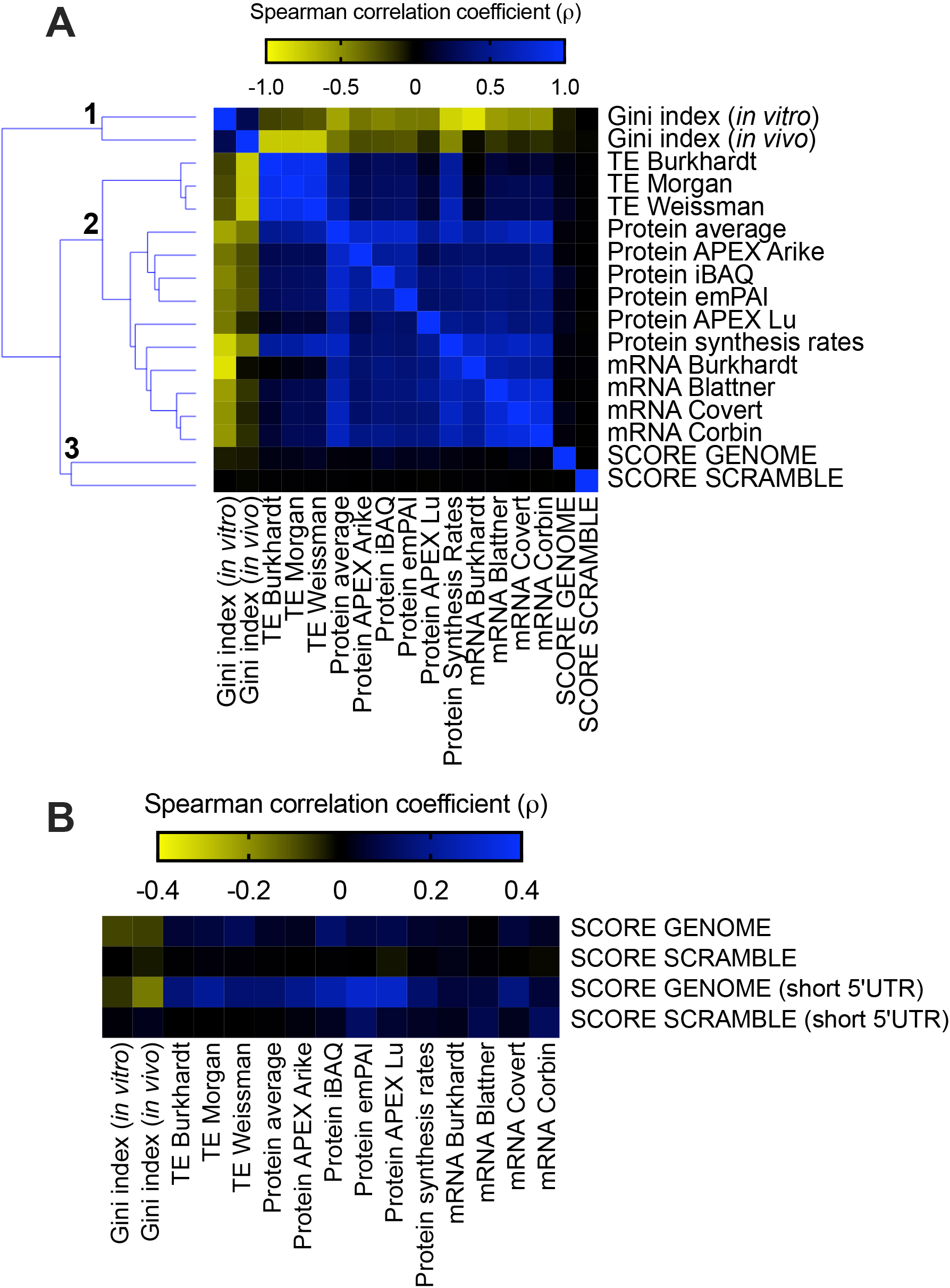
*E. coli* individual gene score calculation and its relationship with different parameters involved with gene expression. (A) Spearman’s correlation between the *E. coli* score derived from the GFP library dataset (SCORE GENOME) with different cellular parameters: Gini index, TE, protein abundance, protein synthesis rate, and mRNA abundance. The heat map shows Spearman’s correlation coefficient (ρ) values ranging from −0.88 (negative correlation, yellow panels) to 1.0 (positive correlation, blue panels). Spearman’s correlation coefficient (ρ) values of the GFP score with other parameters ranging from −0.08 to 0.13. As a control, we used an *E. coli* scrambled genome to calculate the GFP score (SCORE SCRAMBLE). (B) The same analysis described in panel A was performed with a group of genes with short 5’ UTRs (< 25 nucleotides).

This lack of correlation, however, might be due to diverse control mechanisms that operate in a genomic context. It is well known that translation initiation is an important parameter that controls the translation efficiency^24–26^. In bacteria, ribosome binding sites and other regulatory RNA sequences are active control elements for translation initiation^27^. The optimal position and length of the Shine-Dalgarno sequence appeared to be essential for efficient initiation of translation. For example, sequestration of ribosome binding site by mRNA secondary structure negatively affects the initiation of translation^24,27^. To test the influence of this possible regulatory mechanism, we selected genes with short 5’ UTRs (<25 nucleotides) that would be less susceptible to important differences in initiation rates^28^ and repeated the analyses described in panel A of Figure 2. While the results with the SCORE SCRAMBLE data set were barely affected by the selection of genes with short 5’ UTRs, the SCORE GENOME data set exhibited consistently improved Spearman correlation values. Notably, we observed a gain in the correlation coefficients with the RNA structure Gini index (−0.16), TE (0.20) and protein abundance (−0.30) (Figure 2B).

This analysis indicated that the interplay between other determinants of gene expression could significantly skew the expression prediction based on the GFP score. Moreover, one should note that correlation analysis is probably too stringent to detect covariations that are likely not linear. In fact, the GFP score itself does not provide a linear correlation with eGFP fluorescence levels^12^. Therefore, we divided the *E. coli* genes into two groups: one containing genes with high GFP-predicted scores ≥ 4.2 (Figure 3A, green area) and the other with all other genes (< 4.2). The translation efficiency of these two groups of genes was compared using three different datasets^17–19^. Regardless of the TE dataset utilized, the group of genes with higher scores had higher TE when compared to genes with lower scores (Figure 3B-D). The same was true when the protein abundance was compared between low score vs. high score genes (Figure 3E). With this result, we concluded that the sequences related to high GFP expression can enhance the translation efficiency of endogenous genes.

**Figure 3.**
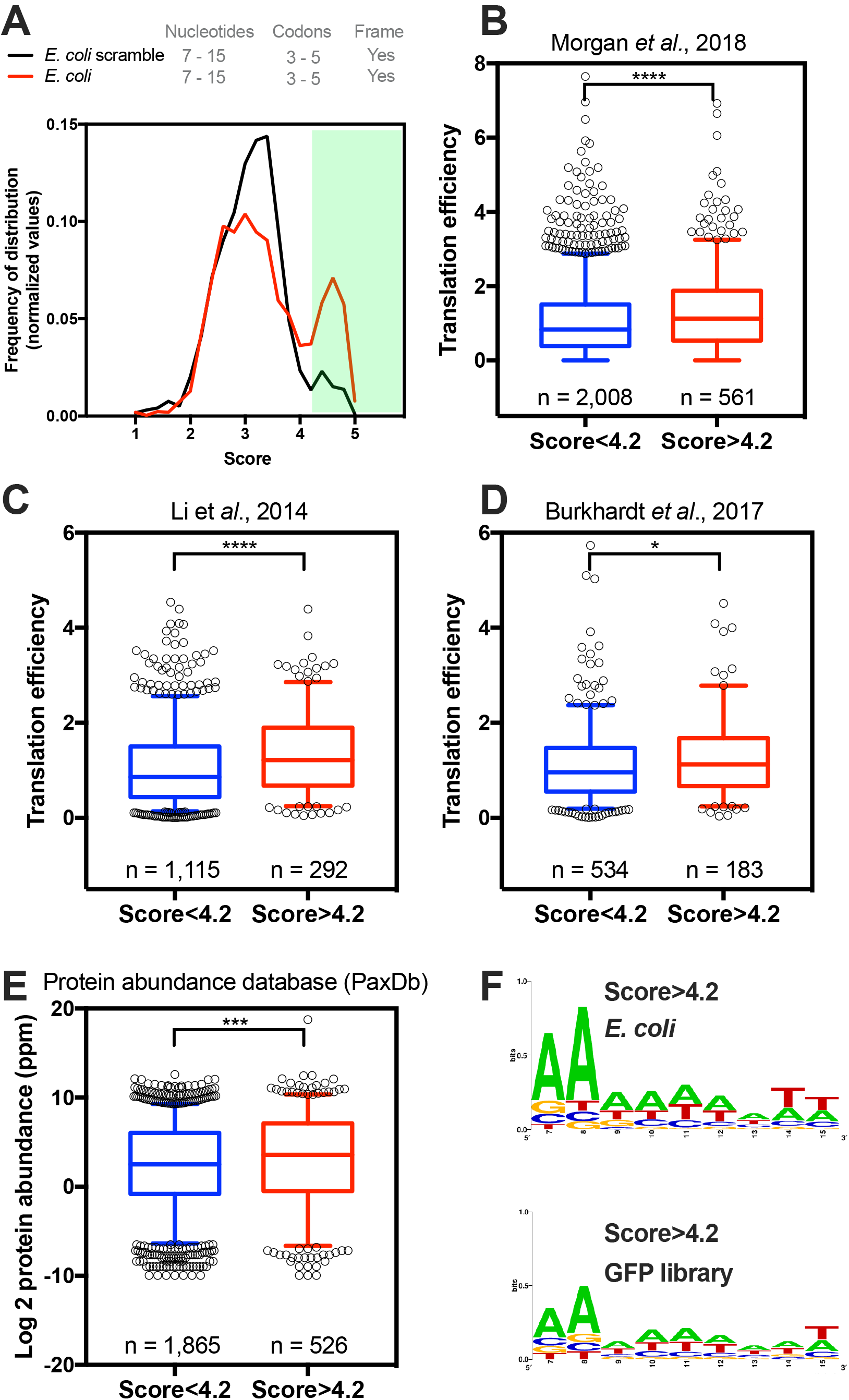
*E. coli* genes with GFP scores higher than 4.2 have higher translation efficiency and protein abundance than other genes. (A) Density histogram of GFP scores for genes identified in *E. coli* for nucleotide composition at codon positions 3-5. As a control, we used a scrambled genome. Based on the GFP score, the *E. coli* genes were divided into two groups: genes with a score higher than 4.2 and scores lower than 4.2. Translation efficiency was measured in three independent studies, Morgan *et al.*, 2018 ^17^(B), Li *et al.*, 2014 ^18^(C) and Burkahardt *et al.*, 2017 ^19^(D). (E) Protein abundance of genes with a score higher than 4.2 and score lower than 4.2. Kolmogorov-Smirnov nonparametric t-test, ****<0.0001, ***0.0004, *0.028. (F) Web Logo of nucleotide composition at codon positions 3-5 of *E. coli* genes or GFP library with a score higher than 4.2.

Using *in vitro* translation experiments, Verma et al. demonstrated that some of the sequences conferring high GFP translatability showed reduced events of ribosome drop-off during early elongation and were more rapidly translated than less efficient sequences^12^. This result apparently conflicts with previous reports arguing that a slow early elongation rate (low-efficiency ramp, or translational ramp) is necessary for high translation efficiency^8^. Such a ramp would prevent ribosome stalling during elongation and subsequent translation abortion events. To check the effect of the sequences associated with high GFP scores on the ribosome early translational rate of endogenous genes, we compared the average ribosome footprint count per nucleotide of the two groups of genes (GFP predicted scores > 4.2 x GFP predicted scores < 4.2). Some ribosome profiling protocols, especially those that include antibiotics, can distort the ribosome profiling data^29–32^, and the best practice for generating ribosome profiling libraries in bacteria is still debated^33^. So, we decided to use ribosome profiling data obtained by two distinct protocols with flash frozen or filtered cells; both protocols avoid the use of translation-interfering antibiotics^33^. We normalized the footprint coverage within the same transcript by dividing each nucleotide read by the sum of the total number of reads for each gene. Independent of the protocol used, the group of genes with higher scores (>4.2) showed less ribosome accumulation at codons 3-5 (nucleotides 7-15) when compared to the genes with lower scores (<4.2) (Figure 4A-B). These data suggest that the translation of the stretch of codons with a high GFP score occurs with higher efficiency and therefore agrees with the experimental data from Verma et al. Gene ontology analysis revealed that genes with higher scores were enriched in cellular component classes related to the cytoplasmic part of the cell (data not shown).

**Figure 4.**
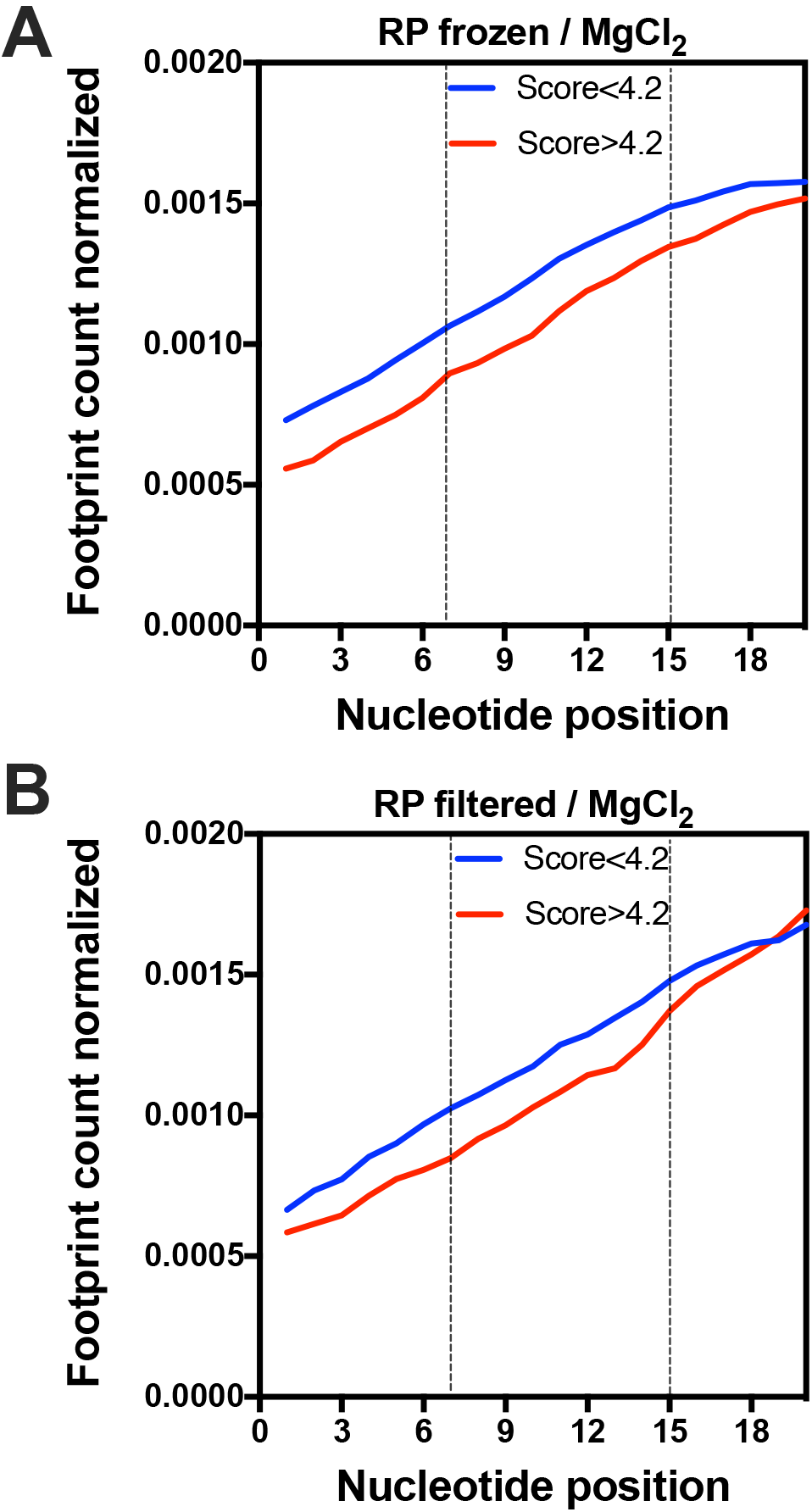
Ribosome occupancy at the first 20 nucleotides of genes with GFP scores higher than 4.2 at codon positions 3-5 (nucleotides 7-15) are lower when compared to other genes. The average ribosome footprint counts of each group were obtained from ribosome profiling (RP) libraries of differently treated samples: frozen/MgCl_2_ (A) or filtered/MgCl_2_ (B)^33^. For each gene, the number of reads per base was normalized to the total number of reads.

The GFP library from Djuranovic’s group contained 259,134 different nucleotide compositions, which is approximately sixty-five times the number of *E. coli* proteins. This means that each *E. coli* gene can have a different nucleotide composition at codons 3-5, and repeated compositions would only exist if some kind of pressure was acting on its selection. Based on this, we asked if the bias towards high GFP-predicted scores was caused by the enrichment of a few conserved high scoring nucleotide sequences or by a multitude of unique sequences. We observed that the vast majority (91%) of the analyzed genes have a unique nucleotide composition at codon positions 3-5 (Figure 5A). Only the remaining 9% of genes possess a repeated nucleotide sequence, where the same sequence is present in at least two different genes (occurrence > 1). This is the same proportion observed with the *E. coli* genes where the GFP predicted score was measured (data not shown). The average GFP predicted score of repeated nucleotide sequences was higher when compared to unique sequences (Figure 5C). To determine whether the higher scores of genes with repeated sequences could reflect a higher efficiency on protein synthesis, we took advantage of ribosome profiling experiment datasets^34^. When the TE of genes with repeated nucleotide sequences was compared to the genome, no statistical difference was observed (Figure 5D). We concluded that the group of genes presenting a GFP-predicted score > 4 was formed by a variety of sequences and not by a few combinations of repeated sequences.

**Figure 5.**
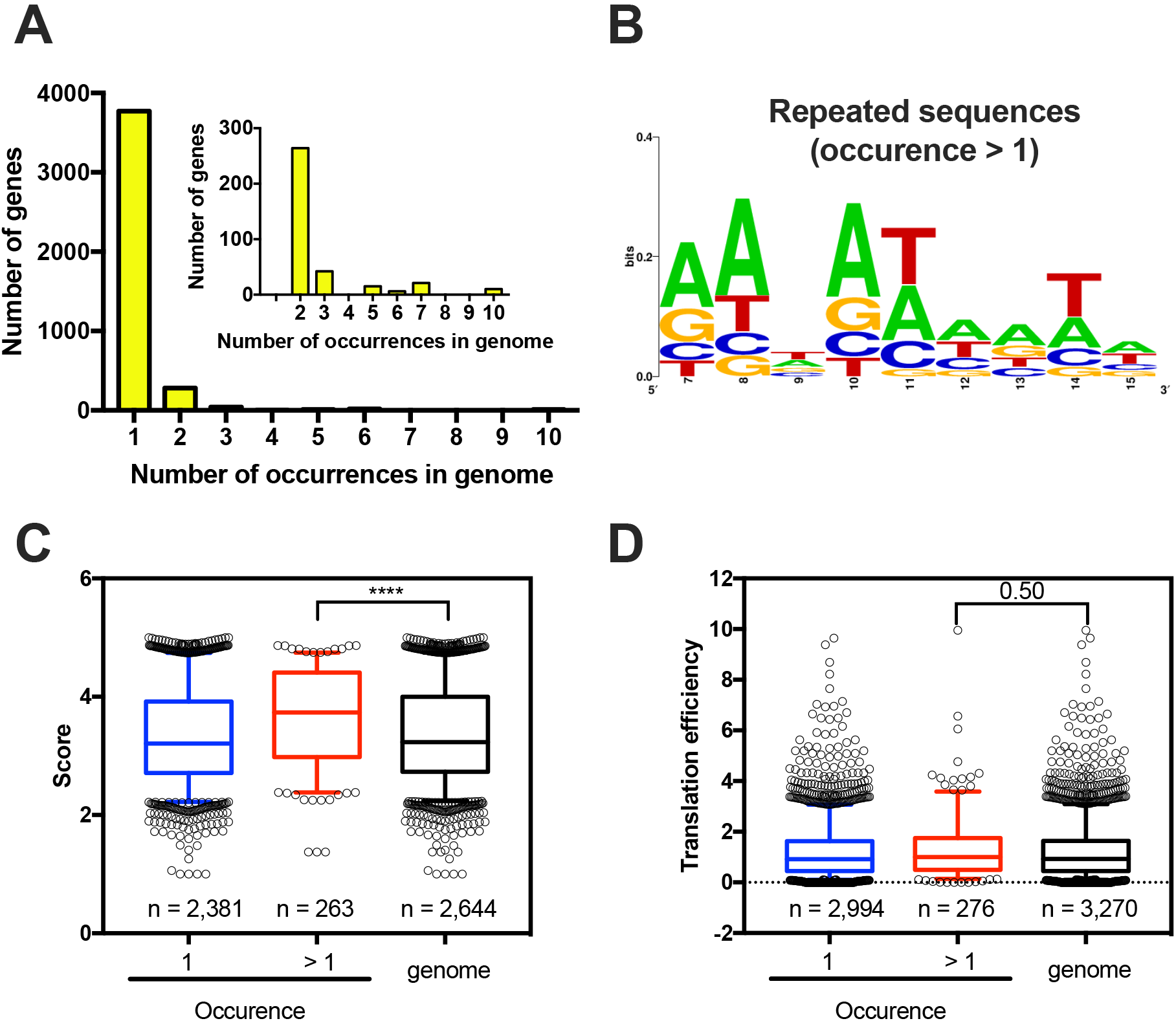
Most *E. coli* genes possess a unique nucleotide composition at codon positions 3-5. (A) The frequency of distribution of nucleotide composition at codon positions 3-5 shows that 91% of *E. coli* genes have a unique nucleotide composition (occurrence = 1). The other 9% share at least one nucleotide composition with another gene (occurrence > 1). (B) Web logo^37^ of genes that share the same nucleotide composition at codon positions 3-5. GFP score average (C) and translation efficiency (D) comparison of genes with a unique nucleotide composition at codon positions 3-5 vs. genes with repeated nucleotide compositions. The TE data were obtained from Morgan *et al.*, 2018^17^. Kruskal-Wallis nonparametric test ****<0.0001, 0.5 = nonsignificant.

Using a motif-scanning approach, Djuranovic’s group found two motifs in GFP variants with scores over 4^12^. The motifs AAVATT (V = A, G or C) or AADTAT (D = G, A or T) are present in several reports at codon positions 3 and 4, yielding higher protein levels than the original construct. We screened the *E. coli* genome searching for these two motifs and found 41 genes (Figure 6A). As a control, we performed the same screening with codon positions 4 and 5, and 34 genes were found (Figure 6A). The 41 genes with one of these two motifs had higher scores when compared to the *E. coli* genome (Figure 6B). Interestingly, genes with these motifs at codon positions 3 and 4 had higher TEs when compared to the genome, but there was no significant difference when the motifs were positioned at codons 4 and 5 (Figure 6C). These data agree with the data that were obtained by Verma et al. through the use of reporters and western blot measurements^12^.

**Figure 6.**
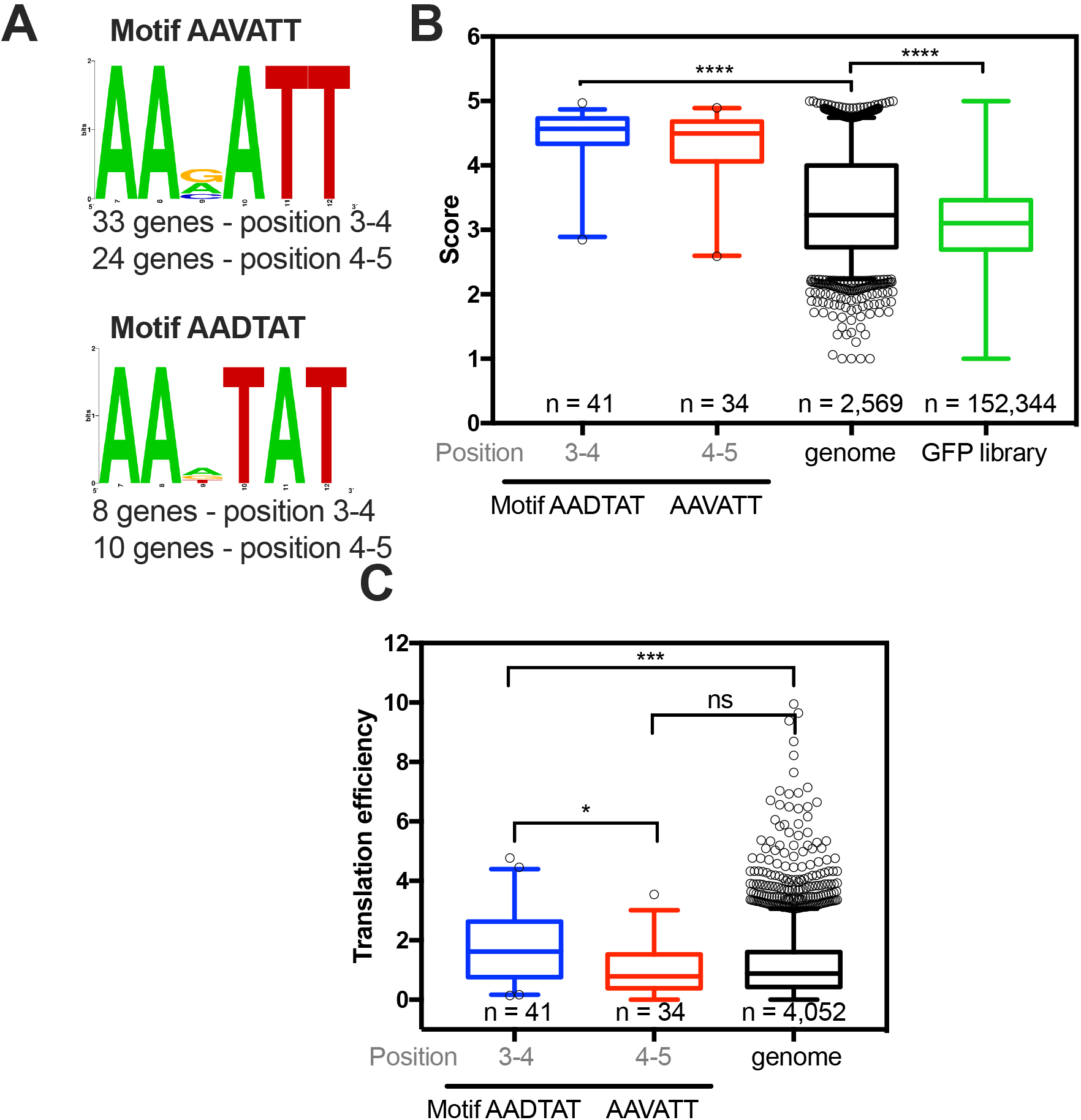
*E. coli* genes with the motifs AAVATT and AADTAT at codon positions 3 and 4 are more efficiently translated than other genes. (A) The motifs AAVATT or AADTAT (V = A, G or C and D = G, A or T) were identified in the *E. coli* genome at codon positions 3 and 4 (33 genes with AAVATT and 8 genes with AADTAT) or at codon positions 4 and 5 (24 genes with AAVATT and 10 genes with AADTAT). The genes with motifs AAVATT or AADTAT at codon positions 3 to 4 have a higher GFP score (B) and translation efficiency^17^(C) than the *E. coli* genome. Kruskal-Wallis nonparametric test *** 0.0005 and ****<0.0001. The TE data were obtained from Morgan *et al.*, 2018.

### Evolutionary conservation of sequences related to high translation efficiency in other bacterial species

To check whether the findings with *E. coli* were valid for other bacterial species, we analyzed three enterobacteria, namely, *Escherichia coli*, *Klebsiella oxytoca*, and *Enterobacter asburiae*. Figure 7A shows that the three analyzed bacteria have a similar bias towards high GFP score values, while the distribution obtained with *H. sapiens* was identical to the GFP library. Each of these bacteria share 1,595 orthologous genes, comprising approximately 40% of each genome. Next, we used this common data set of genes to examine whether the sequences corresponding to codons 3-5 tended to have similar GFP-predicted scores. To exemplify the calculation used herein, we showed the comparison results with three groups of orthologous genes, α, β and γ, corresponding to L-threonine 3-dehydrogenase, sensory histidine kinase QseC and RNA-binding protein YhbY genes, respectively. Differently colored squares represent different nucleotide residues (Figure 7B). A GFP-predicted score was attributed to both windows (3-5 and 9-11), and the results for all 1,595 orthologues were used to build a correlation matrix (Figure 7C). The correlation values obtained with the GFP-predicted scores at codon positions 3-5 were better than those obtained with windows 9-11 (Figure 7C). Additionally, we measured the standard deviation value of the GFP-predicted scores from each group of three orthologous genes and found less variation when the score was calculated at codon positions 3-5 (Figure 7D). For example, 231 groups of orthologous genes had precisely the same score (standard deviation = 0) at codon positions 3-5, while for codon positions 9-10, the number of genes dropped to 151 (Figure 7D, dotted square). We concluded that the translation enhancement effect conferred by certain sequences at codons 3-5 may exert a selection pressure at this particular region of the mRNA.

**Figure 7.**
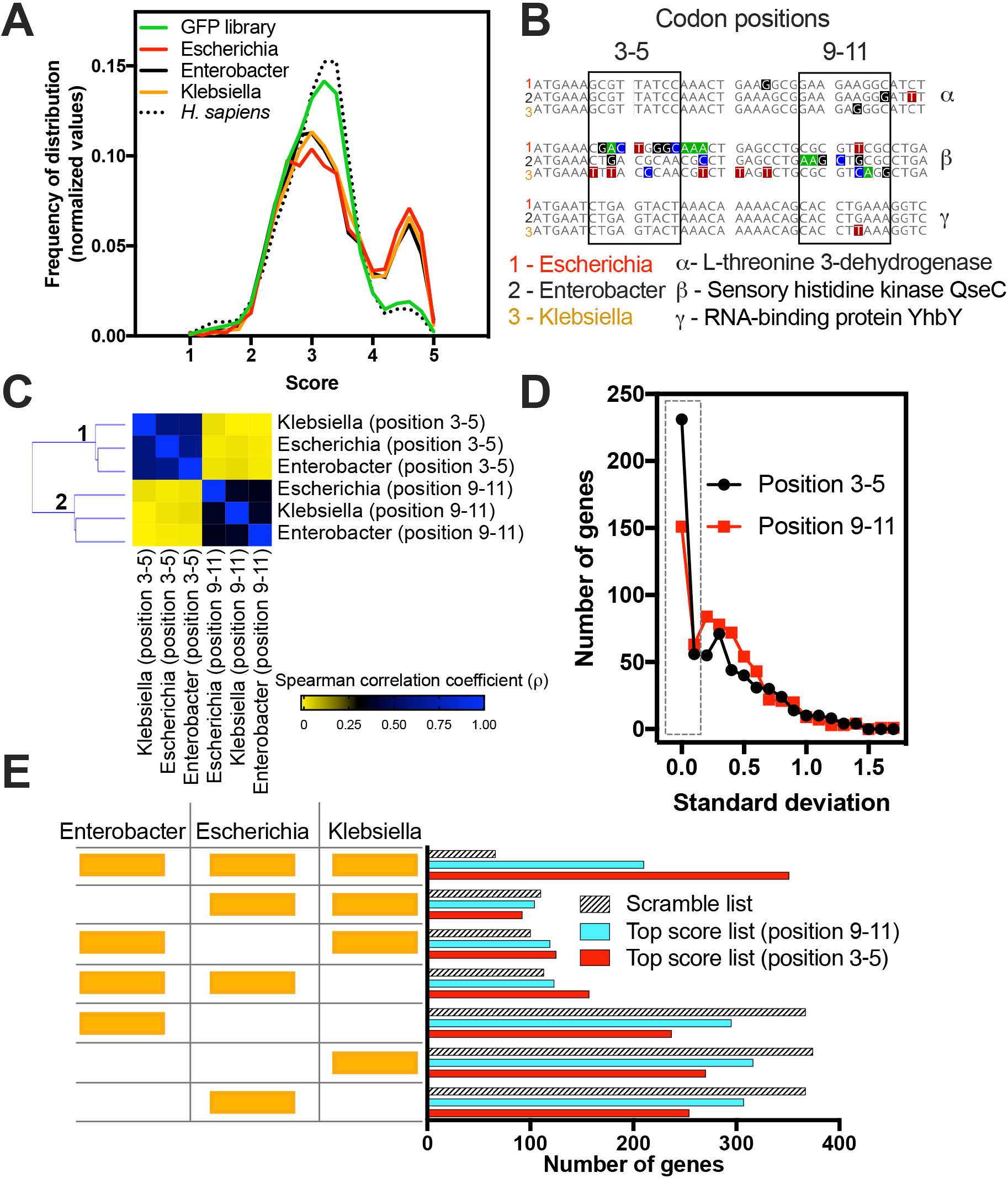
Evolutionary conservation of the short ribosomal ramp. (A) Density histogram of GFP scores of genes identified in *E. coli* (Escherichia), *E. asburiae* (Enterobacter), *K. oxytoca* (Klebsiella) and *Homo sapiens* for nucleotide compositions at codon positions 3-5. As a control, the GFP library score was plotted. (B) A list of 1,595 orthologous genes of *E. coli*, *E. asburiae*, and *K. oxytoca* was analyzed regarding the score of each gene at codon positions 3-5 or 9-11. As an example, three genes (α, β, and γ) are shown. (C) Correlation matrix of the score of orthologous genes of *E. coli*, *E. asburiae*, and *K. oxytoca*. (D) The standard deviation score of each set of three orthologous genes was calculated and plotted as a frequency of distribution. (E) The top 500 genes with the best GFP score of *E. coli*, *E. asburiae,* and *K. oxytoca* were analyzed regarding their orthology. The codon positions 3-5 or 9-11 were used to calculate the GFP score. As a control, a scramble list with 500 genes of each bacterium was used.

To investigate whether orthologous genes are enriched with high GFP scoring sequences, we ranked the total genes of *E. coli*, *K. oxytoca*, and *E. asburiae* in three distinct lists according to the GFP-predicted score obtained for codon positions 3-5. As a control, we created two other lists, one with GFP-predicted scores calculated using position 9-11 and another with scrambled sequences. The top 500 genes with higher scores from each list of the three bacteria (1500 genes in total per analysis) were evaluated in the OrthoVenn2 server, which computes the orthologous gene clusters among multiple species^35^. The number of orthologous genes common to the three species was 351 (23%) for the list obtained with positions 3-5. This was 1.6 times higher than the number of orthologues obtained with the 9-11 list and 5.3-fold higher than the scramble list (Figure 7E, compare the first hatched bar with the first red bar). No important bias was observed for genes restricted to two species.

One possible interpretation for these data is that genes containing sequences associated with high GFP scores demand above average translational efficiency and play essential roles in bacterial physiology; therefore, they tend to be more conserved.

## CONCLUSIONS

Our data suggest an evolutionary pressure selecting some nucleotide compositions at codon positions 3 to 5 since this position affects the TE and protein abundance in a group of enterobacterial genes. On the other hand, our data do not point to a clear correlation between these nucleotide compositions and different cellular parameters, meaning that, at least in physiological conditions, other features of mRNA, such as cis-regulatory elements at the 5’ and 3’ UTRs, codon usage, and mRNA secondary structure, play a more relevant role in translation efficiency. We do not underscore the potential applications driven by Djuranovic’s discovery^12^. The strong effect of some motifs at codon positions 3-5 might have an enormous impact in biotechnology and could be used to improve protein synthesis in recombinant systems. Moreover, the mechanism behind the effect of mRNA composition and protein nascent chain at the beginning of translation impacting ribosome processivity is an open question raised by Djuranovic’s study. It will also be interesting to test in the future the impact of codon composition at positions 3-5 on protein synthesis of some endogenous bacterial genes.

## MATERIALS AND METHODS

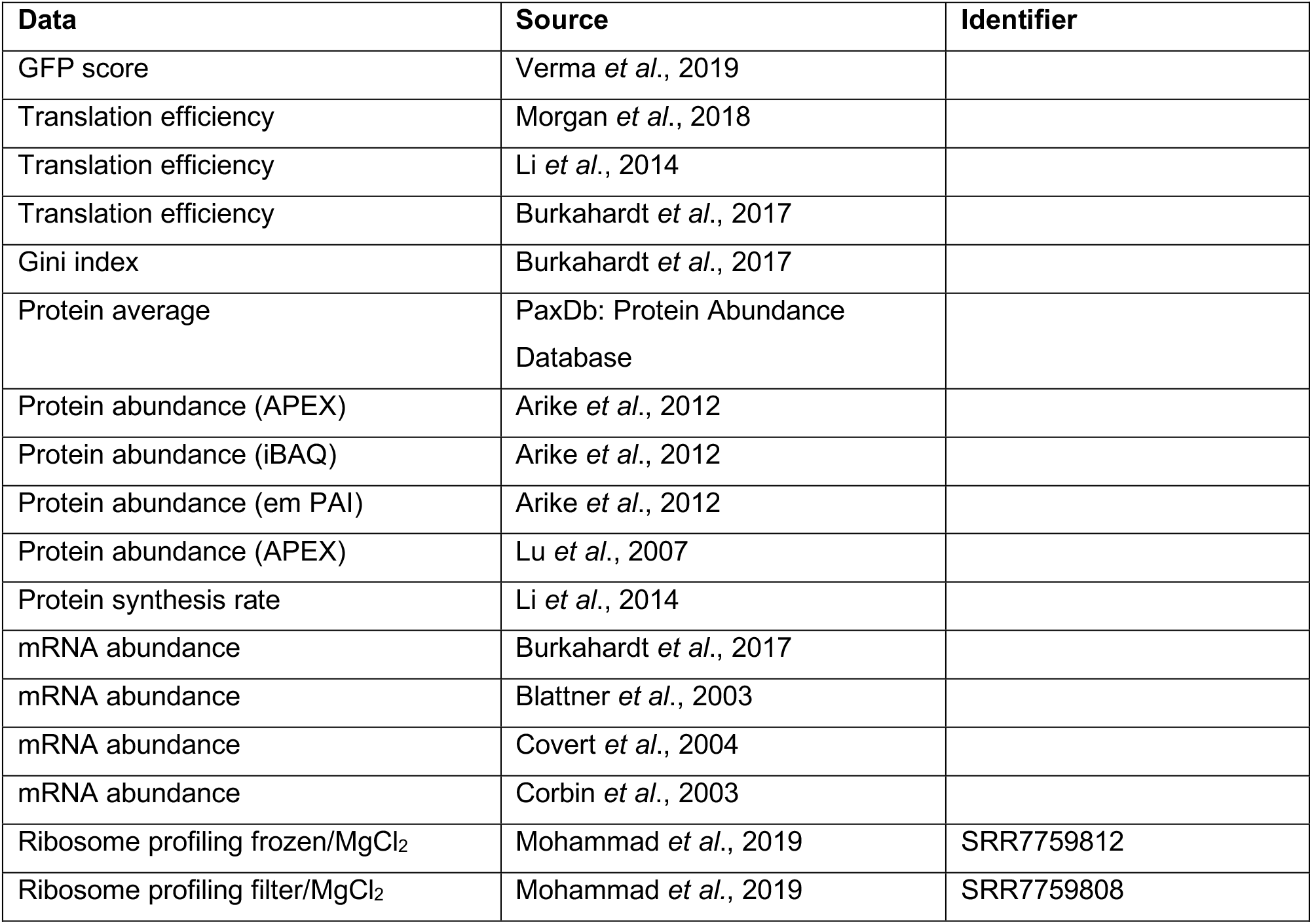
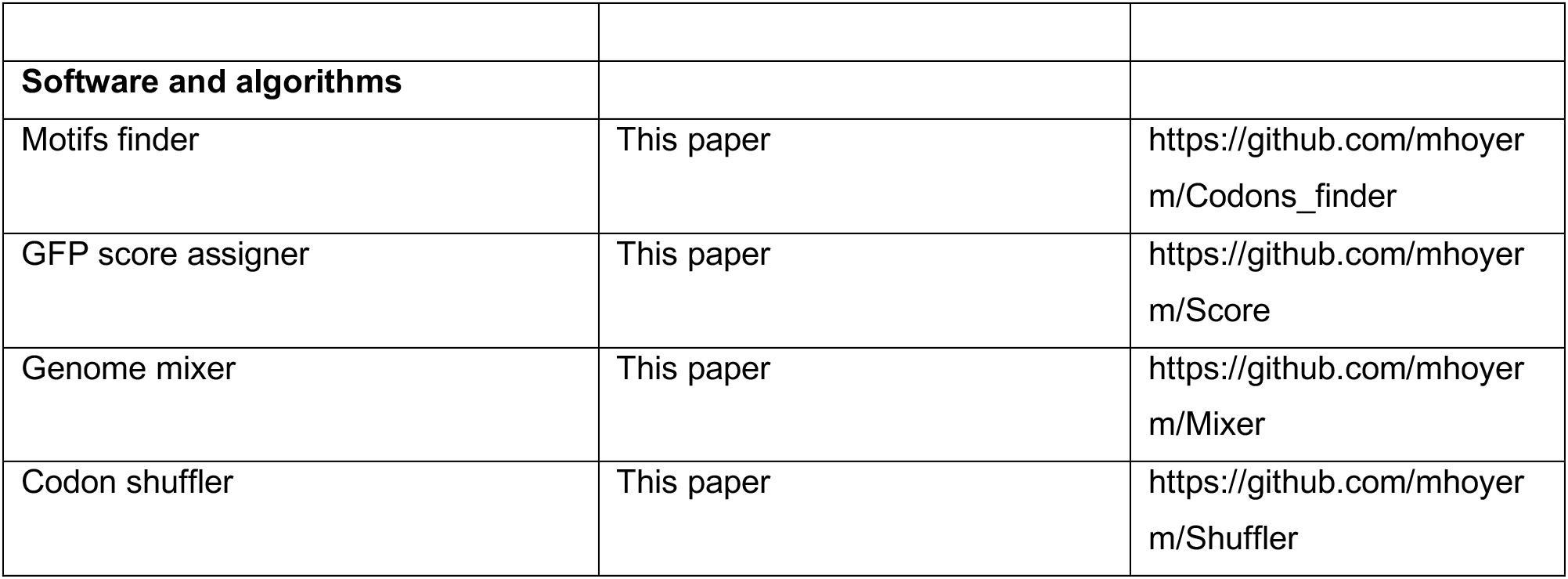

| Data | Source | Identifier |
| --- | --- | --- |
| GFP score | Verma *et al.*, 2019 |  |
| Translation efficiency | Morgan *et al.*, 2018 |  |
| Translation efficiency | Li *et al.*, 2014 |  |
| Translation efficiency | Burkahardt *et al.*, 2017 |  |
| Gini index | Burkahardt *et al.*, 2017 |  |
| Protein average | PaxDb: Protein Abundance Database |  |
| Protein abundance (APEX) | Arike *et al.*, 2012 |  |
| Protein abundance (iBAQ) | Arike *et al.*, 2012 |  |
| Protein abundance (em PAI) | Arike *et al.*, 2012 |  |
| Protein abundance (APEX) | Lu *et al.*, 2007 |  |
| Protein synthesis rate | Li *et al.*, 2014 |  |
| mRNA abundance | Burkahardt *et al.*, 2017 |  |
| mRNA abundance | Blattner *et al.*, 2003 |  |
| mRNA abundance | Covert *et al.*, 2004 |  |
| mRNA abundance | Corbin *et al.*, 2003 |  |
| Ribosome profiling frozen/MgCl<sub>2</sub> | Mohammad *et al.*, 2019 | SRR7759812 |
| Ribosome profiling filter/MgCl<sub>2</sub> | Mohammad *et al.*, 2019 | SRR7759808 |
| **Software and algorithms** |  |  |
| Motifs finder | This paper | https://github.com/mhoyer m/Codons_finder |
| GFP score assigner | This paper | https://github.com/mhoyer m/Score |
| Genome mixer | This paper | https://github.com/mhoyer m/Mixer |
| Codon shuffler | This paper | https://github.com/mhoyer m/Shuffler |

### Data sources

Coding sequences and annotation of *Escherichia coli*, *Homo sapiens*, *Klebsiella oxytoca*, and *Enterobacter asburiae* were obtained from the Ensembl Genome browser (http://ensemblgenomes.org/). The list of *E. coli* genes with short 5’ UTRs was obtained from RegulonDB (http://regulondb.ccg.unam.mx/)^28^.

### Statistical analyses, correlations, and raw data

The raw data used to create Figures 1–4 and for the statistical analyses, including sample size, p-value, D’Agostino & Pearson normality test calculations, Kolmogorov-Smirnov test, when used, are presented in the Supplementary Tables (1-5). For Figure 1A, the best fit equation found was the line equation; the outliers were identified with Q=1% with a confidence level of 95%. All statistical analyses were performed with GraphPad Prism 7 software.

### Gene ontology

The gene ontology analyses were performed in the Gene Ontology Consortium (http://geneontology.org/).

### Motifs finder

In codon positions 3 and 4 or 4 and 5 of an open reading frame genomic sequence, the algorithm searched for sequences of AADTAT or AAVATT, where D is any nucleotide except cytosine, and V is any nucleotide except thymine. When the target sequence was found, the respective gene name and the six nucleotide sequence were written in the output file.

### GFP score assigner

In a defined codon window of an open reading frame from a genomic sequence, the algorithm found the respective GFP score^12^ for the sequence of nine nucleotides and generated an output file containing the gene name, the sequence found, and its respective GFP score.

### Shuffled genome

Data from the original genome were compared to those from a scrambled genome. For that purpose, we designed and implemented two new algorithms that exchanged codons with synonymous codons.

#### Genome mixer

An algorithm that handles the input files by keeping the first codon (ATG) and mixing the positions of the following codons within a genomic sequence, keeping the ratio of nucleotide bases. The random genome generated by this algorithm preserves the number of copies of each codon.

#### Codon shuffler

An algorithm that changes codons in a genomic sequence, maintaining the corresponding amino acid and ratio of nucleotide bases. The random genome generated by this algorithm preserves the number of copies of each codon as well as the amino acid sequence of the original genome.

### Ribosome profiling data

*E. coli* ribosome profiling data were treated as described previously^10,30,34^. The data were analyzed as described by Ingolia and collaborators^34^ except that the program used here was Geneious R11 (Biomatter Ltd., New Zealand) instead of the CASAVA 1.8 pipeline. The data were downloaded from GEO, and the adaptor sequence (CTGTAGGCACCATCAAT) was trimmed. The trimmed FASTA sequences were aligned to *E. coli* ribosomal and noncoding RNA sequences to remove rRNA reads. The unaligned reads were aligned to the *E. coli* genome. First, we removed any reads that mapped to multiple locations. Then, the reads were aligned to the *E. coli* coding sequence database allowing two mismatches per read. We normalized the coverage within the same transcript by dividing each nucleotide read by the sum of the number of reads for each gene.

### Clustering Analysis

All clustering analyses were performed by the Euclidean distance using Orange 3 software^36^.

### Orthologous Analysis

The list of 1595 orthologous genes of *Escherichia coli*, *Klebsiella oxytoca*, and *Enterobacter asburiae* was obtained from OrthoVenn2^35^. The GFP predicted scores at codon positions 3-5 or 9-11 were calculated for each gene. Then, the GFP predicted score correlation of orthologues was measured by a correlation matrix^36^. Additionally, the GFP predicted scores of each group of three orthologues were averaged and the standard deviation was calculated, and the values were plotted as a density histogram (Figure 7D). The scramble list used in Figure 7E was the alphabetical list of genes of each bacterium analyzed. We also used a suit of Orange 3^36^ software to create a scramble list and the same results were obtained (data not shown).

## DATA AVAILABILITY

The algorithms used in this work are available in the GitHub repository (https://github.com/mhoyerm).

## ACKNOWLEDGEMENT

We thank Rafael S. Rocha, Claudio A. Masuda e Maite Vaslin for helpful discussions. We thank Francisco de Oliveira for critically reading the manuscript. We also thank Sergej Djuranovic for helpful discussions and for sharing unpublished data.

**Supplemental data for this article can be accessed on the publisher’s website.**

